# Systemic IgG repertoire as a biomarker for translocating gut microbiota members

**DOI:** 10.1101/2021.08.17.456549

**Authors:** Ivan Vujkovic-Cvijin, Hugh Welles, Connie W.Y. Ha, Lutfi Huq, Shreni Mistry, Jason M. Brenchley, Giorgio Trinchieri, Suzanne Devkota, Yasmine Belkaid

## Abstract

While the microbiota has been associated with diseases states, how specific alterations in composition, function, or localization contribute to pathologies remains unclear. The ability of defined microbes to translocate has been linked to diseases including inflammatory bowel disease (IBD) and was shown to promote responses to immune checkpoint therapy. However, scalable and unbiased tools to uncover microbes with enhanced ability to translocate are limited. Herein, we developed an approach to utilize systemic IgG in an unbiased, culture-independent, and high-throughput fashion as a biomarker to identify gut microbiota members that are capable of translocation across the gastrointestinal barrier. We validate these findings in a cohort of human subjects, and highlight a number of microbial taxa against which elevated IgG responses are unique to subjects with IBD including *Collinsella, Bifidobacterium, Faecalibacterium,* and *Blautia spp*. *Collinsella* and *Bifidobacterium* taxa identified as translocators and targets of immunity in IBD also exhibited heightened bacterial activity and growth rates in Crohn’s disease subjects. Our approach may represent a complementary tool to illuminate privileged interactions between host and its microbiota, and may provide an additional lens by which to uncover microbes linked to disease processes.

**One Sentence Summary:** Circulating microbiota-specific IgG can identify gut microbiota constituents capable of crossing the gut barrier.

## Introduction

Numerous studies have described alterations in the composition of the microbiota in the context of disease states(*1*). However, it is increasingly becoming evident that shifts in relative abundance of microbes may not necessarily imply causality in pathogenesis. Indeed, defined members of the microbiota can initiate critical functions and exert dominant effects on the community irrespective of their relative abundance(*2, 3*). Additional efforts to ascertain the impact of microbiota in disease include metagenomics, meta-transcriptomic and metabolomic studies. However, functional assessment via these approaches remain limited, as nearly half of protein-coding gut bacterial genes have no assigned putative function(*4, 5*) and emergent functions of microbiota gene networks *in vivo* remain poorly understood. Novel, complementary methods to identify microbes with defined behavior or properties would augment biomedical discovery, especially with regards to microbial functions thought to contribute to disease.

Bacterial translocation across mucosal and epithelial barriers has been proposed to contribute to numerous human diseases including inflammatory bowel disease, non-alcoholic steatohepatitis, cancer, HIV-associated inflammation, and obesity(*6–9*). While bacterial translocation is thought to spur pathology among the aforementioned diseases, this phenomenon has also been recently proposed as a mechanism underlying increased responses to immunotherapy(*10, 11*). In other settings, bacterial translocation and/or homing to the tumor environment has been associated with resistance to chemotherapy(*12, 13*). Collectively, these clinical and experimental observations point to bacterial translocation, and ability to engage systemic immunity as a behavior of high relevance to numerous pathological processes in humans. Though significant advances have been made to identify bacteria in extra-mucosal internal organs(*14*), availability of human tissues is limiting and trace environmental contamination can pose a challenge toward identifying microbes in low biomass samples(*15*). On the other hand, the capacity of the adaptive immune system to aggressively respond to microbes encountered beyond barrier surfaces could provide a unique opportunity to identify members of the microbiota able to translocate outside of their physiological niche .

Antibodies to gut microbiota members can be quantified using culture-independent molecular techniques(*16–20*). Such techniques have been successfully utilized to identify mucosal IgA against microbiota members and have facilitated identification of gut mucosal-adherent bacteria able to spur inflammation in several disease models(*16, 17, 20*). Secretory IgA is the predominant immunoglobulin class expressed at mucosal surfaces and contributes to maintenance of mucosal homeostasis via hindering microbial access to epithelia and promoting commensal bacterial colonization, among other functions(*21–25*). In the setting of homeostasis, small intestinal-resident bacteria comprise the majority of gut microbial targets of mucosal IgA, but mucosal IgA can also be preferentially elicited by ingested toxins and gut mucosal-adherent pathogens(*16–18, 20, 26*). Studies quantifying and characterizing IgA-bound members of the microbiota further underscore how immune responses toward defined members can be a better predictor of microbiota localization or function than shifts in relative abundance. How the presence of other classes of antibodies specific to defined members of the microbiota might uncover privileged modes of interaction and localization remains to be addressed.

While IgG responses can often be engaged simultaneously with mucosal IgA against gastrointestinal pathogens, systemic IgG is mounted preferentially during inflammatory challenges(27). Previous work has demonstrated that elevated systemic IgG can be induced in response to translocation of a gut bacterium, and can in that capacity serve to protect from heterologous microbial invasion(28). While gut bacterial taxa can induce systemic IgG at steady state(29), elevated relative levels of IgG against a microbiota member may act as a functional biomarker for identification of microbes with enhanced ability for translocation across the gut barrier in the context of disease states. Whether microbiota member-specific IgG can be probed comprehensively in a high-throughput and unbiased fashion to identify microbes with higher ability to translocate and/or activate the immune system, and whether such a technique can be effective for identifying translocating microbes in humans remain open questions.

Here, we present a method to quantify anti-microbiota IgG titers in human sera/plasma with a surrogate human fecal community from subjects without paired autologous stool, which can readily be applied to already-banked sera from prior human studies. This approach was validated in a cohort of Crohn’s disease and ulcerative colitis patients. Thus, assessment of anti-microbiota IgG repertoires may represent a powerful approach to identify microbiota members able to translocate, a property that may contribute to defined immune or pathological states.

## Results

We first assessed whether systemic IgG targets different bacteria than those targeted by mucosal IgA. To this end, a cohort of human subjects was profiled for IgG and IgA reactivities to autologous fecal microbiome samples using magnetic bead-based separation and 16S rRNA profiling of immunoglobulin-bound and unbound fecal bacteria (“IgA-seq” and “IgG-seq”, respectively). IgG and IgA scores were calculated as log-ratios of taxon abundances in the bound and unbound fractions (**Fig. 1A-B**). While concordance was evident between IgG and IgA scores, several gut bacterial taxa exhibited greater than 10-fold differences in IgG scores compared to IgA scores (**Fig. 1C**) indicating differential targeting of taxa by these two immunoglobulin classes. Notably, among these differentially-targeted taxa, IgA scores correlated more strongly with baseline fecal relative abundances than did IgG scores (**Fig. 1D-E**). Thus, systemic IgG reactivity against gut bacteria differs from that of mucosal IgA and is less strongly linked to fecal bacterial relative abundance.

**Figure 1:**
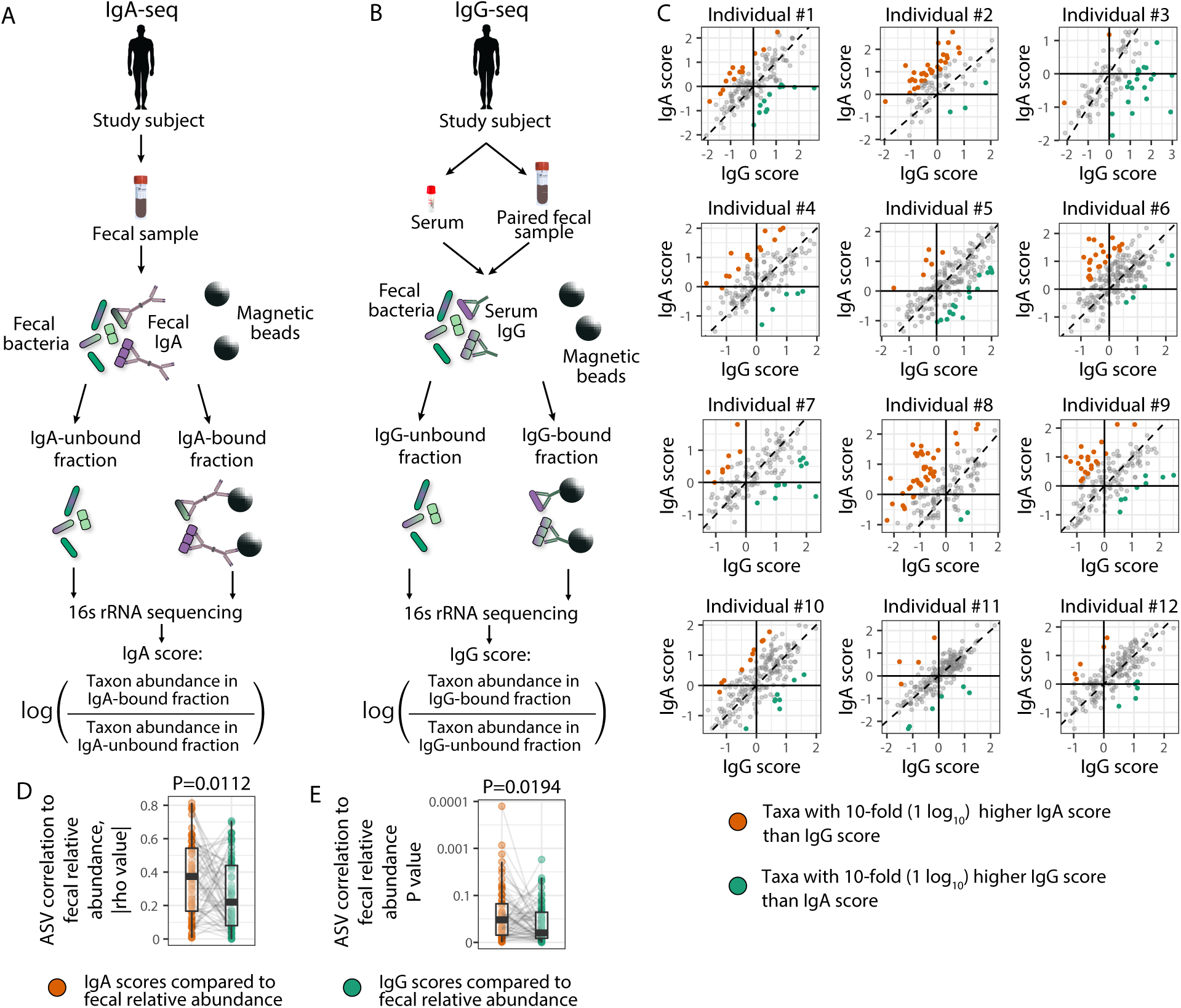
Fecal IgA and systemic IgG target different gut microbiota constituents. A) IgA-seq workflow. Fecal bacteria from mice and humans were incubated with phycoerythrin (PE)-conjugated anti-IgA antibodies, after which anti-PE microbeads facilitated magnetic separation of IgA-bound bacteria from IgA-unbound bacteria. These two fractions were subjected to 16S rRNA sequencing and IgA scores for each taxon were calculated as the log-ratio of taxon relative abundance in the IgA-bound fraction over that in the unbound fraction. B) IgG-seq workflow. Fecal bacteria were incubated with autologous sera and stained with PE anti-IgG antibodies, fractionated, and sequenced as for IgA-seq. C) Numerous gut bacterial taxa exhibit differential binding by IgA and IgG within individuals. Taxa with ten-fold higher IgA scores than IgG scores are depicted in orange, while taxa with 10-fold higher IgG scores are in green. D-E) IgA scores correlate more strongly with baseline fecal relative abundances than IgG scores. D) Shown are absolute values of all Spearman rho values for correlations comparing IgA or IgG scores to fecal relative abundances among taxa whose IgA and IgG scores differed by ten-fold (colored points in panel C). E) Spearman P values for the same comparisons as in (D). Paired ratio t-tests were performed for (D) and on log-transformed values of (E).

To better understand the genesis of this heterogeneous bacterial-specific targeting by systemic IgG, we sought to test the hypothesis that systemic IgG reactivity is enriched in specificities directed toward translocating bacteria. To test this specific point we utilized *Toxoplasma gondii,* a well-described model of mucosal infection associate with both microbial translocation and increased reactivity toward the microbiota(*30, 31*). To identify viable microbiota constituents that preferentially translocated across the gut barrier during *T. gondii* infection, mice were orally infected and peripheral organs (liver, spleen) were subjected to bacterial culture and identification. A separate group of mice were infected with *T. gondii* and the immunoglobulin repertoires were allowed to form over 4 weeks, after which IgA- and IgG-seq were performed (**Fig. 2A**). As expected, confirmed translocator bacteria (those detected in peripheral organs among two or more independent experiments) exhibited significant differences in IgG levels within mice experiencing *T. gondii*-induced barrier breach as compared to uninfected mice (**Fig. 2B-C**). This observation was not evident when examining mucosal IgA (**Fig. 2B,D**), emphasizing the possible unique utility of elevated IgG as a biomarker for translocating microbes.

**Figure 2:**
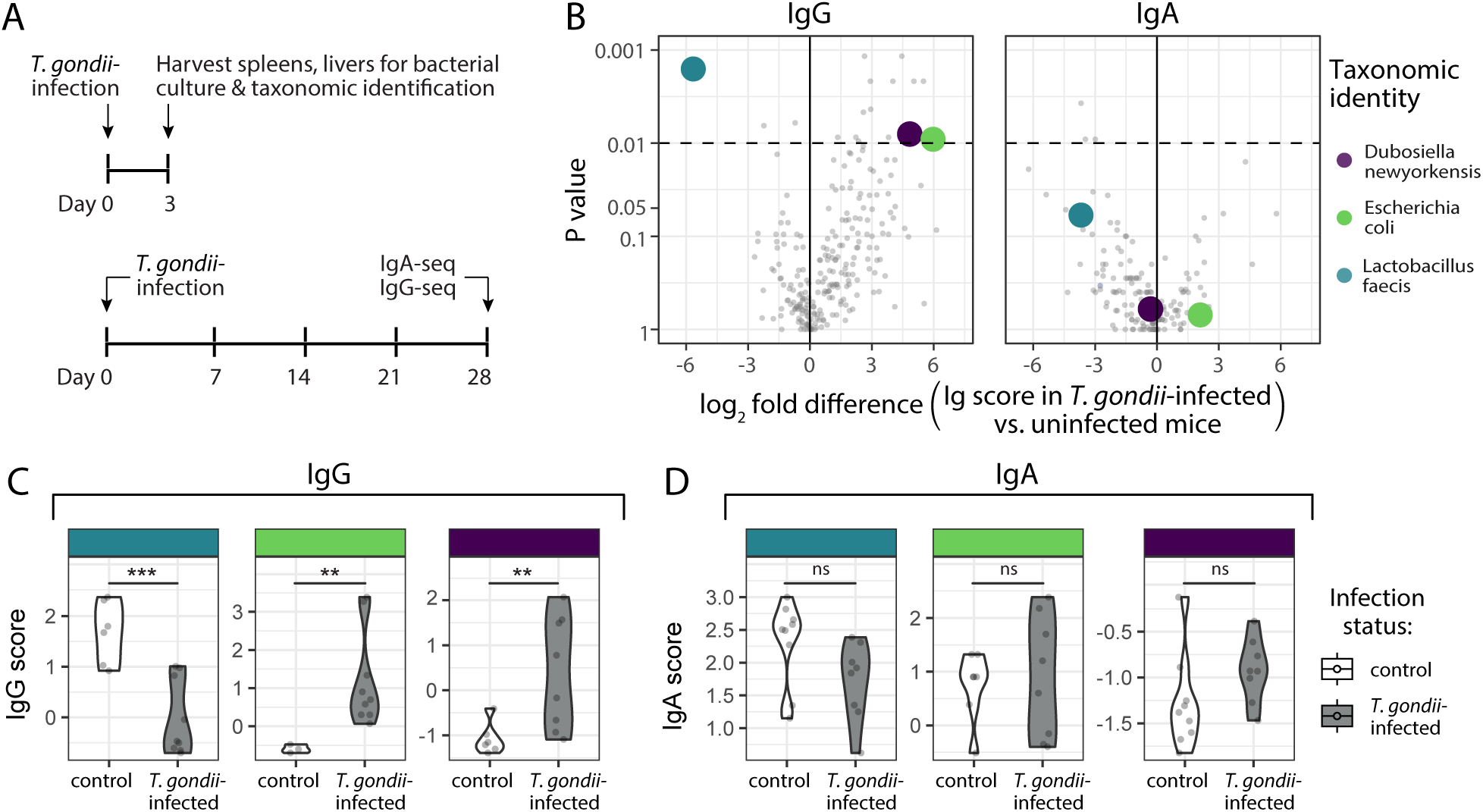
Systemic IgG identifies invasive gut bacteria capable of translocation and inflammation. A) Schematic of experimental plan. Livers and spleens of acute T. gondii-infected mice were processed and subjected to bacterial culture at day 3 post-infection. Colonies were identified using full-length 16S rRNA sequencing. IgA-seq and IgG-seq were performed on n=17 mice at day 28 post-infection. B) Shown are translocators that were cultured from livers or spleens in two or more independent experiments. Mean fold differences in IgA and IgG scores between T. gondii-infected and uninfected mice were calculated for each taxon, and Mann-Whitney U tests were performed to calculate P values. C) IgA score dynamics are shown for translocator taxa highlighted in (B). D) IgG score dynamics are shown for translocator taxa highlighted in (B). *** P<0.005; ** P<0.01.

Mice and humans exhibit substantial differences in gut microbiota composition, and salient immunomodulatory gut bacteria of mice are absent in humans(*32, 33*), highlighting the importance of developing scalable research tools that can be used in humans using readily available sample sources. Availability of paired serum and stool is limited within the majority of prior human studies performed to date. To overcome these important hurdles, we developed a modified IgG-seq procedure using human subject sera in conjunction with a surrogate fecal community (SFC) to be used in place of autologous, paired stool (**Fig. 3A**). This SFC was designed to contain the greatest number of unique bacterial taxa possible from a combination of stool samples derived from healthy donors, to ensure probing the greatest number of taxa for serum IgG reactivity. This was accomplished by first profiling forty healthy human volunteer stool samples via 16S rRNA sequencing. The combination of stool samples that yielded the community with maximal diversity was subsequently calculated using Monte Carlo-based *in silico* modeling (**Fig. S1A**). Fifteen healthy human volunteer stool samples were thus combined and used as the SFC for subsequent analyses. The resulting SFC encompassed 8 phyla, 13 classes, 17 orders, 33 families, 84 genera, and 1,876 unique amplicon sequence variants of human gut bacteria (**Fig. S1B**). When comparing IgG scores using the SFC (SFC-IgG-seq) to IgG scores obtained using paired serum and stool (IgG-seq) on the same cohort, there was significant, non-random concordance between the two methods (**Fig. S1C-D**).

**Figure 3:**
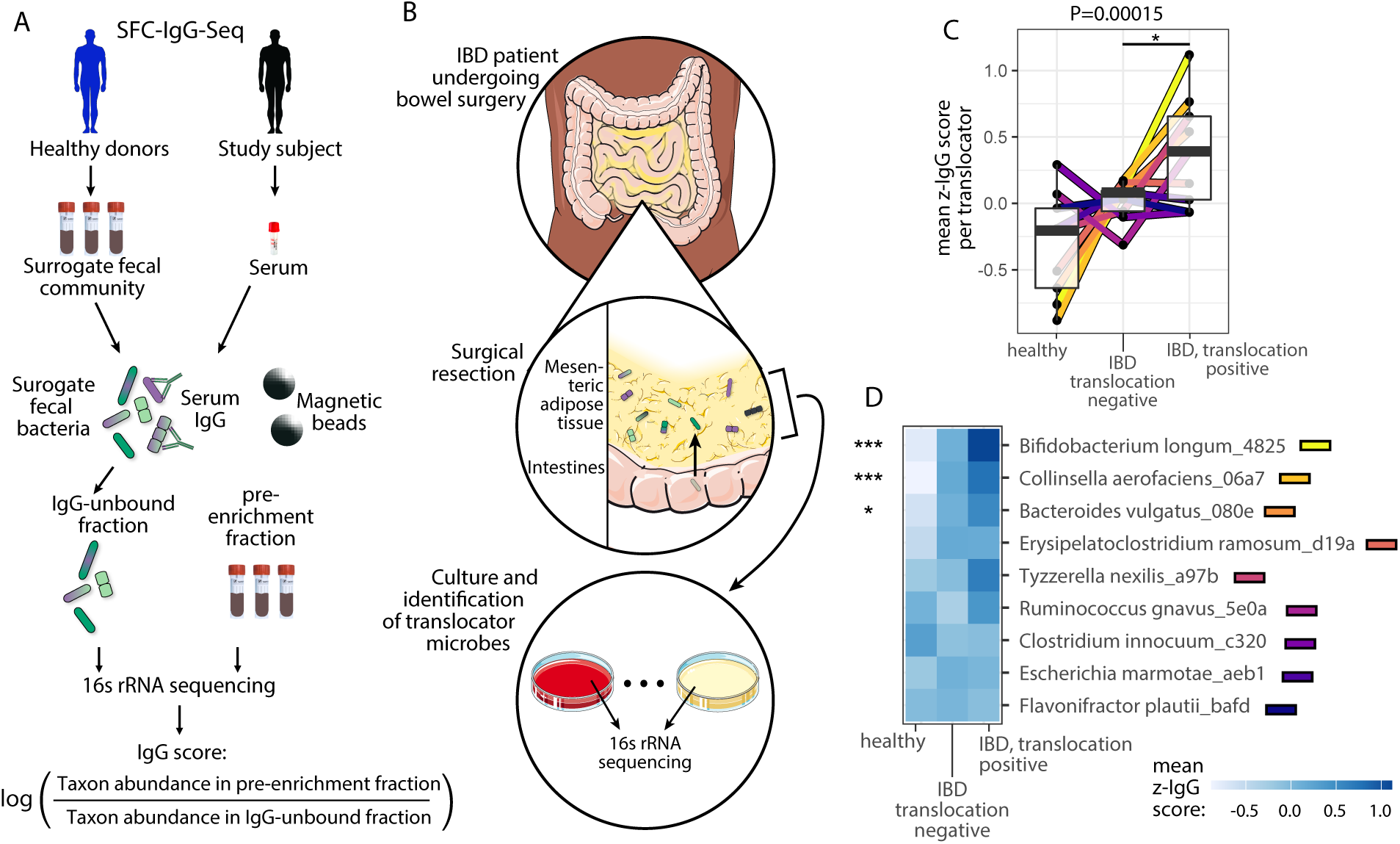
Translocator bacteria exhibit heightened IgG scores in humans. A) SFC-IgG-Seq workflow used on sera of subjects in panel (B). In absence of stool from study subjects, a surrogate fecal community (SFC) comprising a combination of healthy donor feces from a separate cohort was engineered (see Supp. Fig.). Fecal bacteria and sera are co-incubated and the IgG-unbound fraction is sequenced along with triplicate pre-enrichment aliquots of the SFC. IgG scores are calculated comparing taxon abundances to the pre-enrichment community (see Supp. Fig.). B) Workflow for identification of translocating bacteria in a cohort of IBD patients undergoing medically indicated bowel resection surgery. Mesenteric adipose tissue from resected small and large intestinal samples was subjected to bacterial culture, and full-length 16S rRNA sequencing was performed on colonies to identify translocator bacteria on a per-individual basis. C) Translocator bacteria exhibit heightened IgG scores in IBD subjects with confirmed translocation. For each taxon, mean z-IgG scores were compared within healthy, IBD subjects with no detected translocation for that taxon, and IBD subjects for whom that taxon was cultured from the mesenteric adipose (translocation positive). These groups were treated as ordinal variables in that order and linear regression on IgG scores was performed grouping all taxa (P=0.00015). IgG scores for translocator bacteria were also elevated in IBD translocation-positive subjects as compared to IBD translocation negative (P=0.0268, paired two-sided Student’s T-test). D) Mean z-IgG scores are shown for each taxon within each subject grouping. Linear regression was performed on each individual taxon, with significance values summarized at the left. Raw IgG scores were converted to row z-scores for analysis and visualization. *** P<0.001, *P<0.05. IBD, inflammatory bowel disease; SFC, surrogate fecal community.

To understand whether systemic IgG may be used as a biomarker to facilitate identification of translocating gut bacteria, we applied SFC-IgG-seq to a cohort of inflammatory bowel disease subjects that underwent ileal or colonic surgical resections and healthy controls (cohort characteristics in **Table S1A-B**). Mesenteric adipose tissue from the IBD subjects was subjected to bacterial culture as a way to identify taxa that had translocated across the gut barrier and entered the mesentery(*34*) as depicted in **Fig. 3B**. Recovered isolates were identified by full-length 16S rRNA sequencing as previously reported(*34*) and translocator bacteria that were isolated from mesenteric adipose tissue of four or more subjects were considered for downstream analyses. We sought to test whether the translocation of such bacteria in a given individual was associated with heightened cognate systemic IgG responses. We compared IgG scores in IBD subjects for which translocation was confirmed via culture to IgG scores in subjects for whom translocation was not observed, on a per-translocator basis. When aggregating translocator bacteria, a significant increase in IgG scores was seen in translocation-positive subjects as compared to translocation-negative subjects (P=0.0268, **Fig. 3C**). Given that IBD disease exhibits dynamic and fluctuating severity within individuals over time, it is possible that the time of mesenteric sampling missed some translocation events. Thus, we analyzed IgG score dynamics for translocator bacteria among a third control group of healthy subjects. We observed a significant stepwise increase in IgG scores across these healthy subjects, translocation-negative IBD subjects, and translocation-positive IBD subjects (P=0.00015, **Fig. 3C**) when considering all translocator bacteria. Individual translocator bacteria that exhibited significant linear increases in IgG scores from healthy, to IBD translocation-negative, to IBD translocation-positive subjects included *Bifidobacterium longum, Collinsella aerofaciens,* and *Bacteroides vulgatus* (**Fig. 3D**). To understand whether mucosal abundance alone explained heightened IgG scores for detected taxa, we examined correlations between IgG scores and mucosal relative abundances as detected by 16S rRNA profiling of mucosal biopsies of the same cohort of individuals. Such correlations were generally weak and exhibited a random distribution (**Fig. S2A**), supporting the idea that IgG scores are non-redundant with gut mucosal relative abundance. Small intestinal residency of a gut bacterium may be sufficient to induce mucosal IgA(*18*). Thus, we examined whether presence of each taxon at the mucosa, assessed by 16S rRNA gene sequencing of intestinal biopsies, impacted observations of heightened systemic IgG in translocation-positive human subjects. When examining only subjects with detectable mucosal presence of a given translocator, the observation that translocator bacteria exhibited heightened IgG scores in IBD subjects with confirmed translocation as compared to IBD subjects without translocation remained evident (P=0.0142, **Fig. S2B**).

IBD, which encompasses the two disease classifications of Crohn’s disease (CD) and ulcerative colitis (UC), is characterized by barrier breach and immune activation. This immune activation is a critical driver of disease progression and is thought to be stimulated and amplified by the gut microbiota. Proportions of fecal bacteria that are endogenously coated with IgG are heightened in both CD and UC(*35–39*) and this measure correlates strongly with disease activity(*40, 41*). However, the specific identities of IgG-targeted bacteria in IBD and what such IgG-targeting of a gut microbiota constituent may indicate with regards to its function remain poorly understood. Consistent with prior observations(*35–41*), our analysis revealed that a markedly greater proportion of bacteria in the surrogate fecal community were targeted by IBD subject sera than by sera of healthy controls (**Fig. 4A**). Using SFC-IgG-seq, we examined IgG-targeting of all detectable taxa within the SFC, first comparing total anti-microbiota IgG repertoires between subject groups. IBD subject anti-microbiota IgG repertoires significantly clustered separately from healthy healthy subjects (**Fig. S3**), a finding that was robust for both CD and UC separately (**Fig. 4B-C**). This segregation by anti-microbiota IgG repertoires was independent of concurrent medications when examining all subjects (**Table S1C**). Numerous specific taxa were found to be preferentially targeted by IgG in CD and UC subjects as compared to healthy healthy controls (**Fig. 4D-F**). Systemic IgG responses mounted against microbial antigens can have cross-reactivity against structurally related antigens such as those from phylogenetically related taxa(*28*). Such off-target responses are typically weaker than those directed at the immunogen(*42, 43*), and thus taxa with greatest group-wise differences in IgG scores are more likely to be true targets of IgG. Such taxa include *Bifidobacterium longum, Collinsella aerofaciens, Faecalibacterium prausnitzii,* and *Blautia wexlerae* (**Fig. 4G-J**). Anti-microbiota IgG repertoires segregated CD subjects from controls more strongly than UC subjects from controls (F=2.121 vs. F=1.78, Fig. **4B-C**). In line with this observation, the majority of taxa that differed in their IgG scores between CD and UC subjects exhibited heightened IgG scores in CD subjects (**Fig. 4K**), suggesting heightened targeting of gut microbiota members in CD as compared to UC.

**Figure 4:**
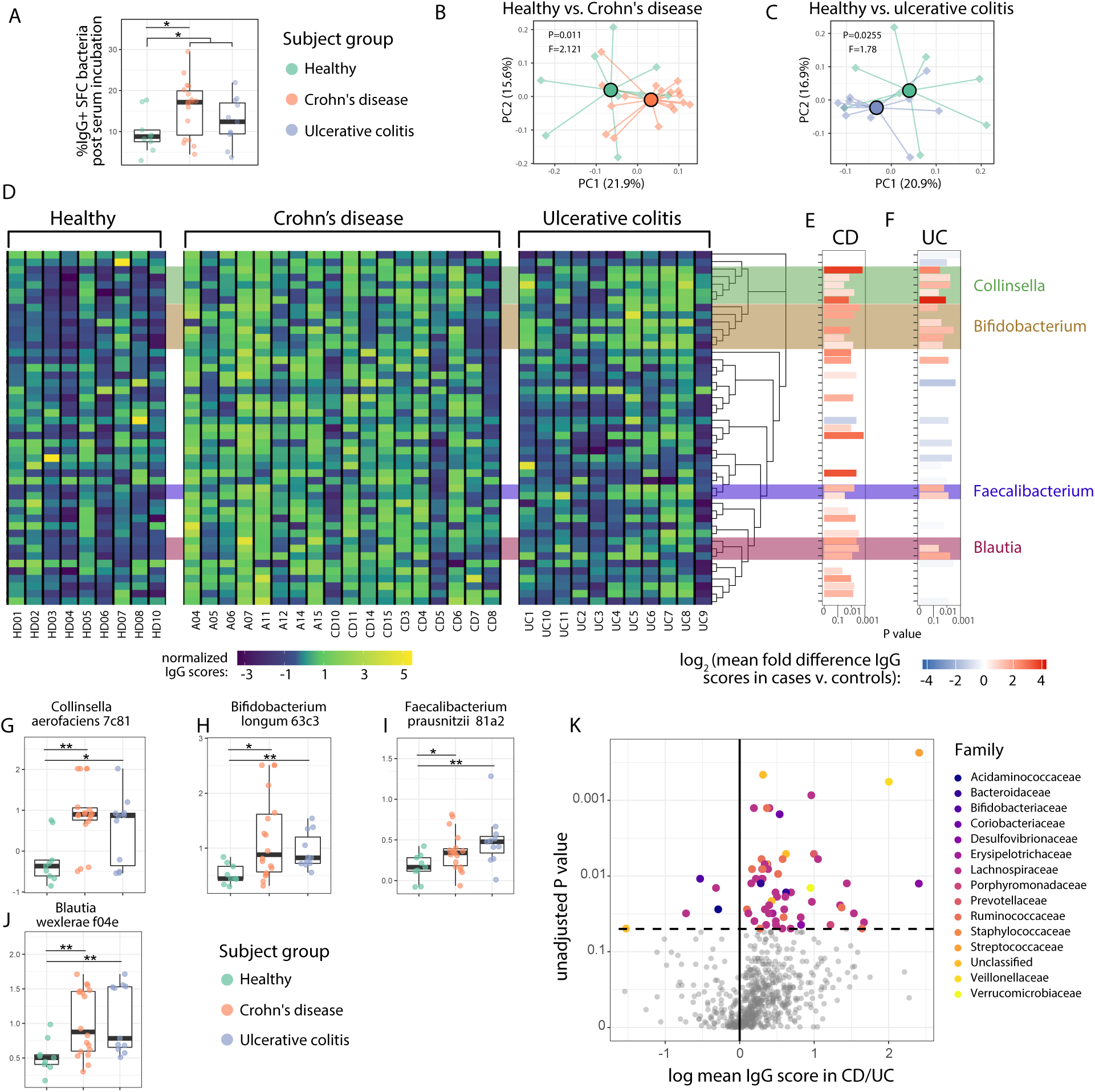
Anti-microbiota IgG profiles differ significantly in IBD vs. healthy human subjects. A) Percentages of bacteria in the SFC bound by IgG were determined for each subject via flow cytometry. Healthy vs. Crohn’s P=0.015, healthy vs. IBD P=0.029, healthy vs. ulcerative colitis P=0.26; two-sided Student’s T test. B-C) Principal coordinates analyses using pairwise Pearson correlation coefficient matrices encompassing IgG scores for all taxa. P values and F statistics were calculated using PERMANOVA. D) Heatmap of all taxa whose IgG scores differed among the three subject groups (Kruskal-Wallis unadjusted P<0.02). E-F) Taxa whose IgG scores differed significantly between healthy and Crohn’s disease subjects (E), and between healthy and ulcerative colitis subjects (F), by Mann-Whitney U test P<0.05. G-K) Plots depicting IgG scores of representative members of genera that differed in IgG scores between healthy and IBD subjects. Shown are IgG scores for *Collinsella aerofaciens* (G), *Bifodobacterium longum* (H), *Faecalibacterium prausnitzii* (I), and *Blautia wexlerae* (J). K) IgG scores in CD and UC subjects for all taxa were compared using Mann-Whitney U tests, with resulting unadjusted P values on the y axis (log-transformed) and log fold differences in CD vs. UC subjects along the y axis.

Efforts to characterize bacterial functions in human microbiome studies include the assessment of human gut commensal bacterial growth rates via meta-transcriptomics and metagenomics. Such studies have revealed differences in proliferative activity of specific taxa in healthy subjects as compared to a number of diseases(*44*), though the behavior of bacteria that exhibit differential growth patterns/activity and whether such bacteria engage in privileged interactions with their host remains poorly understood. Focus was placed on CD given greater IgG profile differences, and data were examined from the longitudinal integrative Human Microbiome Project (iHMP) consortium(*45*). Taxa were ranked by the strength of correlation between the extent of dysbiosis in CD subjects and overall transcriptional rate (RNA/DNA ratio) of each taxon (**Fig. 5A**), a measure linked to bacterial activity(*46*) and/or growth rate(*47*). Taxa whose transcriptional rate correlated with dysbiosis overlapped with taxa identified via IgG-seq (in **Fig. 4D**) and/or identified as translocators via cultivation (in **Fig. 3B**). Specifically, in the iHMP dataset we examined taxa that we confirmed as translocators in our cohort (highlighted in green, **Fig. 5**), taxa that exhibited heightened IgG scores in our CD subjects (purple, **Fig. 5**), as well as taxa that were both confirmed translocators and exhibited heightened IgG (red, **Fig. 5**). Several taxa that fell within these three categories were represented among the top 10 taxa whose transcriptional rate (RNA/DNA ratio) was most correlated with dysbiosis in the iHMP (**Fig. 5A**).

**Figure 5:**
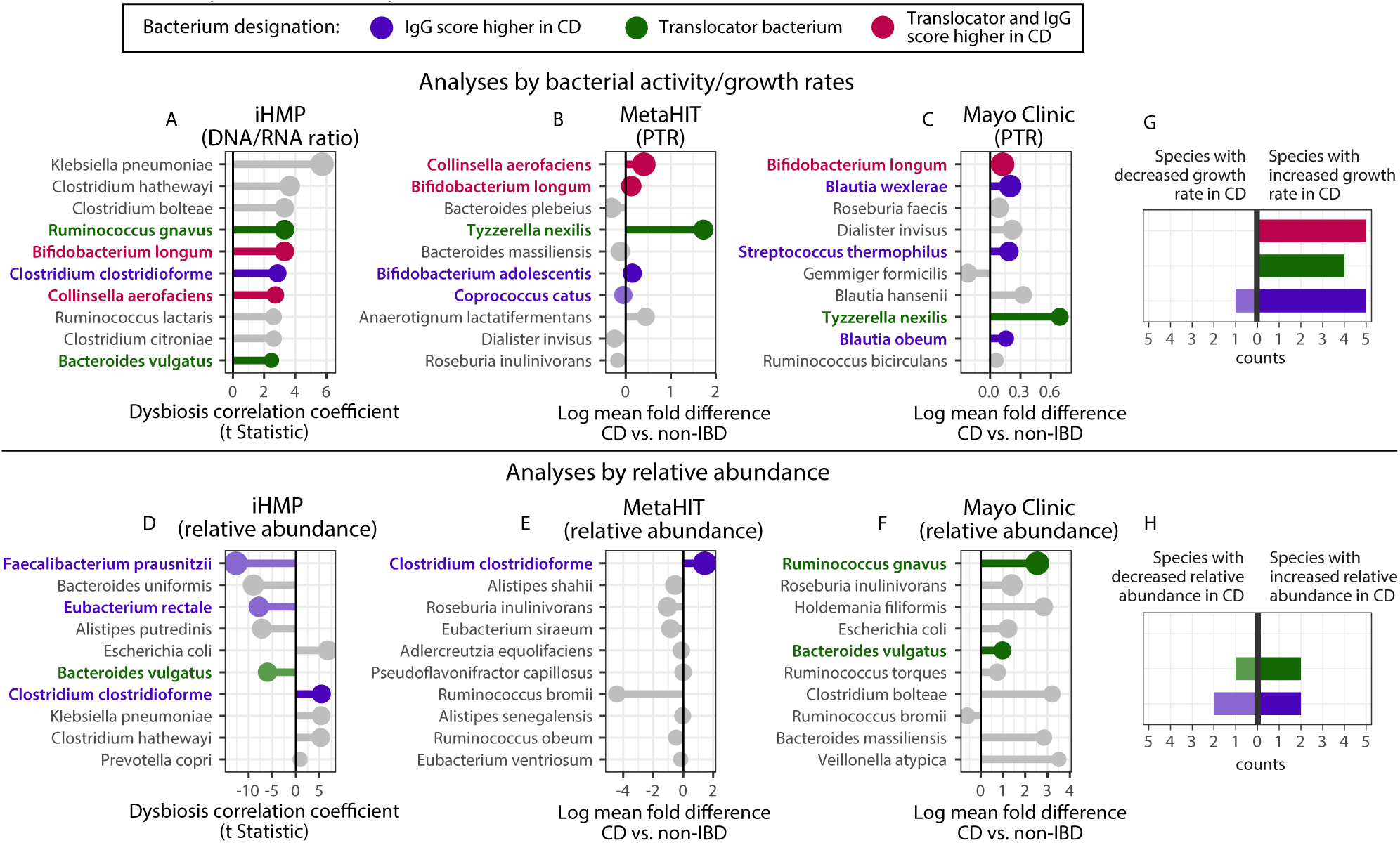
Bacterial growth dynamics correlate more strongly with translocator status and IgG score dynamics in Crohn’s disease than does relative abundance. A-F) Analyses of external, independent metagenomics datasets of Crohn’s disease (CD) and healthy subjects, by markers of bacterial growth (A-C) and relative abundance (D-F). Species were ranked by resulting P values and are shown in order with most significant at the top of each graph. The top ten taxa classified at the species level for each analysis are shown, with P values in Supp Table 2. Bacteria of the following categories are highlighted in color: those with higher IgG scores in CD (as defined in Methods), those that were cultured from >4 mesenteric adipose tissue subject samples in our IBD cohort, those that were both cultured from mesenteric adipose tissue and exhibited heightened IgG scores in CD. A) RNA/DNA ratios for each bacterium, a proxy for bacterial cell growth, were compared as described previously(*1*) to a marker of longitudinal dysbiosis within CD subjects, and the resulting strength of correlation is shown on the x-axis. B-C) Peak-to-trough ratios (PTR), also a proxy for bacterial cell growth, were calculated and compared between CD and healthy subjects using Mann-Whitney U tests. Species were ranked by resulting P values. D-F) Analyses of the same datasets as in (A-C), though examining relative abundance of species instead. D) Relative abundances were compared to a marker of longitudinal dysbiosis within CD subjects, with the resulting strength of correlation is shown on the y-axis. E-F) Relative abundances were compared between CD and healthy subjects using Mann-Whitney U tests as in (B-C). G) Count-based summary of (A-C), adding all numbers of each bacterial designation: translocators (green), IgG-targeted species (purple), or both IgG-targeted and translocators(red). Counts of species are separated by direction of trend, as either being enriched or depleted in CD. H) Count summary as in (G) but for relative abundances (panels D-F).

Bacterial growth rates can be inferred from metagenomic sequencing studies via the quantification of peak-to-trough ratios (PTR), which measure the ratio of reads near a bacterial origin of replication compared to reads at the replication terminus(*44, 48*). In two independent cohorts of CD and healthy subjects from separate continents(*49, 50*), we compared bacterial growth rates of all detected gut taxa between CD and healthy subjects. Concordant with the analysis of bacterial activity by transcriptional rate, several taxa which were translocators and/or exhibited heightened IgG in our cohort of CD subjects overlapped with those exhibiting increased bacterial growth rates assessed by PTR (**Fig. 5B-C**). Intriguingly, *Bifidobacterium longum*, which was a prominent translocator and target of systemic IgG in our IBD cohort, exhibited higher bacterial activity/growth rates in all three independent datasets queried (**Fig. 5A-C**). *Collinsella aerofaciens* was also a prominent translocator and target of systemic IgG, and showed higher activity/growth rates in two of three datasets (**Fig. 5A-B**). *Tyzzerella nexilis,* a confirmed translocator in our cohort, was also among the top species exhibiting higher growth rates in two of three datasets (**Fig. 5B-C**).

In comparison to analyses examining differential bacterial growth rates in CD vs. healthy subjects, relative abundance analyses yielded lists of differentially abundant taxa that had comparatively less representation of translocators and bacterial targets of IgG (**Fig. 5D-F**). Taxa that were confirmed translocators and/or exhibited heightened systemic IgG in CD were rarely identified by differential abundance analyses (7 instances), and exhibited mixed directional results with nearly equal numbers of taxa being enriched in CD as enriched in healthy controls (**Fig. 5H**). On the other hand, when examining the top taxa with differences in bacterial growth rates between CD and healthy controls, bacteria that were translocators and/or exhibited higher IgG scores were represented in 15 instances, with 14 out of 15 exhibiting uniformly higher bacterial growth rates in CD (**Fig. 5G**). Thus, translocators and pro-inflammatory gut bacteria in CD were more represented amongst the taxa identified by differential bacterial growth rate analyses than those by relative abundance analyses (P=0.032, chi-square test) when comparing CD to controls. In aggregate, these data indicate that heightened bacterial activity/growth rate in CD is a trait that overlaps with heightened bacterium-specific IgG responses and the functional capacity of microbes to translocate.

## Discussion

Here, we report that the enhanced ability of gut commensal bacteria to elicit heightened systemic IgG responses upon translocating across the gastrointestinal tract can be leveraged to identify translocating bacteria in the context of human disease. We find that in both UC and CD, uniquely heightened systemic IgG responses are detectable against several gut taxa including those within the *Bifidobacterium, Collinsella, Faecalibacterium,* and *Blautia* genera, suggesting preferential translocation of these taxa in IBD. As bacterial translocation in immunocompetent hosts is met with immune activation, the principal driver of IBD pathology, this microbial activity is likely relevant to disease progression and thus translocator bacteria may represent a viable therapeutic target. CD subjects exhibited a greater number of taxa targeted by IgG and greater differences from control subjects with regards to their anti-microbiota IgG repertoire. Taxa with heightened IgG responses in CD subjects and/or those detected as translocators in CD also exhibited greater activity and proliferation across three independent metagenomic and meta-transcriptomic datasets, suggesting a link between increased overall bacterial activity and the capacity to translocate.

While *Collinsella* and *Bifidobacterium spp.* have not exhibited consistent relative abundance-based differences between CD and healthy subjects in prior human microbiome association studies, other prior evidence suggests a possible role for these bacteria in disease pathways. Indeed, high *Collinsella* and *Bifidobacterium spp.* abundance has been associated with positive response to anti-TNF therapies, suggesting a link between these bacteria and this pathologic innate inflammatory pathway(*51*). *Bifidobacterium spp.* have also been shown to cause small intestinal inflammation and pathology(*52*), and to induce large and small intestinal T_H_17 cells (*53, 54*), a cell type implicated in human inflammatory bowel disease(*55*) as well as murine models(*56*).

Not all translocator bacteria in our study exhibited comparative elevation of bacterium-specific IgG in subjects for which translocation was confirmed. It is possible that such gut bacteria have evolved immune evasion strategies(*57*), or that they may translocate over the lifetime of a healthy individual to a sufficient extent as to induce a plateau of maximal specific IgG in both cases and controls. The observation of IgG against a bacterium also does not necessarily denote translocation in every case. Indeed, previous work demonstrated that a small fraction of the microbiota can be targeted by IgG independently of confirmed translocation as in the case of *Akkermansia muciniphila*(*29*). Approaches such as those which we describe that include control subjects without presumed translocation are likely to help discern steady-state IgG production against a bacterium from that of IgG mounted in response to bacterial translocation. Furthermore, our population of IBD subjects exhibited severe manifestations of disease requiring surgical interventions. Thus, examination of IBD subjects across a spectrum of disease severity will inform whether gradations of IgG titers against the specific taxa identified here may manifest differentially across disease severity and putative sub-types.

While the IgG-seq technique as utilized herein will preferentially highlight gut bacteria that are differentially targeted by high-avidity IgG responses, low affinity, T cell-independent IgA and IgG that are broadly reactive to the majority of gut commensal microbes have been observed during homeostasis(*18, 19*). Given that induction of such homeostatic responses does not require T cells, such responses are likely induced by qualitatively different host-microbe interactions than those highlighted in the present study. Group comparisons (e.g. IBD vs. healthy) allow discrimination of bacteria associated with IgG responses that likely surpass the threshold for homeostatic antibody induction. Examination of different antibody isotypes and multimerization states may facilitate finer discrimination of different modes of host-microbial interaction. For example, serum IgA, which differs structurally from secretory IgA expressed at the mucosa, is robustly induced by systemic inflammatory stimuli(*58, 59*) like serum IgG and can also protect from lethal bacteremia(*60*). Further investigation into the role of secretory vs. serum IgA is warranted and may provide another lens by which to understand host-microbe immune relationships. Finally, in the context of systemic immune engagement with a translocating microbe, cross-specific antibody induction may occur and contribute to elevation of IgG against structurally related antigens such as those from phylogenetically related taxa. In studies examining such processes, systemic antibodies exhibit greater propensity for binding the primary immunogen than to heterologous antigens(*61, 62*). Thus, we placed emphasis on the top taxa exhibiting differential IgG scores between cases and controls in expectation that the true immunogen would have increased IgG-targeting and that taxa targeted by non-specific cross-reactive antibodies would have comparatively smaller differences in IgG scores between cases and controls. Given the intentionally designed microbial complexity of the SFC, it is likely the original immunogen is present in this community and thus the relevant IgG targets can be identified. However, the immunogen may be expressed by an especially rare taxon or one absent from the SFC and thus possible cross-reactivity without detection of the original immunogen cannot be ruled out.

The method developed in the present study to identify bacteria targeted by systemic IgG yielded information that was non-redundant with that of relative abundance comparisons, providing an additional tool in the pursuit of identifying causally-related microbiota constituents in disease. Specifically, quantifying IgG against gut microbiota members using the IgG-seq technique as presented herein facilitated identification of gut bacteria that harbored the discrete function of having translocated across the gut barrier. Comparing IgG levels against all gut bacteria between healthy and IBD subjects revealed a repertoire of anti-microbiota immune responses that was unique to IBD, highlighting potential new bacterial targets in this disease The SFC-IgG-seq technique using a surrogate fecal community as presented can be readily used on existing serum sample sets to identify translocator gut microbiota members in human inflammatory disease, without need for stool sampling. Utilizing a single combined stool substrate per subject also obviates variability in comparative gut microbiome studies that is inherent to human stool, including intra-individual temporal variability and intra-fecal spatial variability(*63*). Thus, examining immune responses to a fixed surrogate fecal community can reduce noise attributable to certain sources of variability in human microbiome studies. Mining host immune memory to better understand microbiota member function and behavior as presented may help illuminate therapeutic targets for the amelioration of immunopathology and/or immune adjuvants in the setting of cancer.

## Materials and Methods

### Study Design

Murine and human study sample numbers are reported in sections dedicated to each experimental design. Mice were male C57BL/6J-CD45a(Ly5a) background mice obtained from Taconic Biosciences, and each experiment utilized mice of the same age. Murine experiments were each performed on three independent occasions. Each SFC-IgG-seq subject sample was assayed in duplicate; two wells of SFC were incubated with human subject plasma and were processed as described below.

Human sera for SFC-IgG-seq were randomly distributed throughout the 96-well plate so as to minimize possible sources of bias. The initial healthy human cohort (n=12) with paired serum and stool was used for discovery purposes to examine differences between anti-microbiota IgA and IgG profiles and was obtained from the CEPHIA consortium of the University of California – San Francisco. The subsequent human cohort (n=42) with only serum was used to address the hypothesis that translocator bacteria could be identified by their IgG titers and included healthy, Crohn’s disease, and ulcerative colitis patients from Cedars-Sinai Medical Center. All human subjects provided informed consented for participation in research. Blinding was not performed and investigators were aware of translocation metadata for samples, so as to perform the analyses described.

### IgA- and IgG-seq on paired human serum and stool samples

Stools and sera from 12 healthy human donors who provided informed consent were obtained and IgA-seq and IgG-seq were performed simultaneously. Sera were heat-inactivated at 56C for 35 minutes to block complement activity and were diluted to 1:500. 200 mg of stool were weighed on dry ice and resuspended in 1% bovine serum albumin (w/v) in phosphate buffered saline, a solution used for all subsequent staining, washing, and resuspension steps. Suspensions were centrifuged at 200 rcf for 1 minute to pellet food particles. Each sample input was adjusted to have equivalent OD_600_, and 5 separate aliquots were taken: 2 to be incubated with serum for IgG-seq, 2 for mucosal IgA-seq (without serum incubation), and one for pre-enrichment 16S rRNA sequencing. One aliquot from each pair of aliquots was used for isotype staining while the other was used for downstream fractionation and analyses. Aliquots for IgG staining were incubated with paired sera from the same individual for 30 minutes at 4C. Samples were washed, with all centrifugation steps being performed at 8,000 rcf for 3 minutes. Serum-incubated samples were stained with anti-IgG PE (Miltenyi #130-093-193, 1:50 dilution) and SYTO-62 (Invitrogen, 1:500 dilution) which stains bacterial cells, and samples for IgA-seq were stained with anti-IgA PE (Miltenyi #130-093-128, 1:40 dilution) for 20 minutes at 4C. Aliquots of each stained sample were taken for flow cytometry assessment of quality of staining.

Samples were then subjected to magnetic column-based separation using anti-PE microbeads (Miltenyi, #130-048-801) at a concentration of 1:20, and subsequent steps were followed as per manufacturer protocol and the Ig- eluate was collected and spun by centrifugation to remove excess buffer prior to freezing. To increase degree of Ig+ cell enrichment, the Ig+ fraction was subjected to an additional column enrichment. The column used for the first enrichment was washed by passing ethanol through the column with a plunger followed by PBS, and the Ig+ sample was then applied and the manufacturer protocol was again performed. The final Ig+ fraction was centrifuged, excess buffer was removed, and was frozen for downstream processing.

DNA was extracted from samples using the MagAttract PowerMicrobiome DNA/RNA Kit (Qiagen). Amplification of the V4 16S rRNA gene (primers 515f, 806r) was performed in accordance with the Illumina recommended protocol, amplicons were purified using AMPure XP beads (Beckman Coulter), were quantified using the KAPA Quantification Kit (Roche), and pooled at equimolar concentrations. Pooled libraries were sequenced on a MiSeq instrument (Illumina).

Sequencing analysis was performed as follows. Amplicon sequence variant (ASV) tables were constructed using Dada2(*64*) in the R programming environment. ASV tables were rarefied at 11,000 sequences per sample. IgA and IgG scores were calculated as the log_10_ ratio counts of each taxon in the Ig-enriched microbial fraction over the counts in the Ig-depleted microbial fraction. Scores for each ASV were compared to baseline pre-enrichment ASV relative abundances across all individuals using Spearman correlation tests (R package ‘Hmisc’).

### Toxoplasma gondii infections

An ME-49 clone of Toxoplasma gondii transfected with red fluorescent protein was obtained from Dr. Michael Grigg (NIAID/NIH), which was used for cyst production in wildtype C57BL/6 mice and for cyst quantification via fluorescence microscopy. Tissue cysts were obtained from orally infected mice and resuspended in phosphate buffered saline. Experimental mice were infected with 8 cysts via oral gavage. All animal experiments were performed in accordance with guidelines from the National Institutes of Health Animal Care & Use Committee.

### Translocator identification from culturing internal murine organs

Spleens, livers, and brains from *T. gondii*-infected mice were removed using autoclaved forceps and scissors, and were transferred to pre-reduced anaerobic transport media tubes (Anaerobe Systems, #AS-911) prior to transfer to an anaerobic chamber. Tissues were subsequently physically disrupted and filtered through sterile 40 micron filters using pre-reduced anaerobic phosphate buffered saline. Resulting suspensions were plated on YCFAC+B (Anaerobe Systems, #AS-677), MTGE (Anaerobe Systems, #AS-777), and Columbia Blood Agar (ThermoFisher, #R01217) at 37C for 48-72 hours. A set of Columbia Blood Agar plates was also incubated in aerobic conditions at 37C for 48-72 hours. Brain tissue was cultured as a negative control, as were buffers in which tools were dipped prior to organ resection, both of which routinely yielded no colonies. Colonies were propagated in liquid media corresponding to that from which they were grown, were cryopreserved, and were identified by full-length 16S rRNA Sanger sequencing. Translocator culture was performed during three independent experiments with 5-10 mice infected with *T. gondii* being subjected to culture each time, with 1-2 uninfected control mice to test for negative culture plates. Culture of translocator bacteria from mesenteric adipose tissue of human subjects undergoing surgical resection was performed as described previously(*34*). Human translocating bacterial isolates were identified by Sanger sequencing of full-length 16S rRNA amplicons.

### IgG- and IgA-seq on *T. gondii*-infected mice

Sera and stool were collected from mice 4 weeks post-infection with *T. gondii* and frozen as part of three independent experiments with 5-10 mice per group (infected and uninfected). Stools were placed in buffer (1% bovine serum albumin in phosphate buffered saline, used for all subsequent washing and staining steps), were physically disrupted using sterile pipette tips, and were homogenized by repeated pipetting using wide-bore pipette tips by pipetting. Stool suspensions were spun at 50 rcf for 1 minute to pellet food particles. Supernatant was transferred via 40 micron filter to a new tube, and washed by centrifugation at 8,000 rcf for 3 minutes (same speed and time for subsequent washes). 1/10^th^ of this suspension was used to quantify OD_600_ for normalization of input samples. Another 1/10^th^ of this suspension was set aside for flow cytometry-based assessment of pre-enrichment IgA and IgG fecal bacterial binding. Samples were split for IgA- and IgG-seq. Samples for IgG-seq were incubated in 1:50 plasma for 30 minutes at 4C while samples for IgA-seq rested at 4C. Samples were resuspended in stain mixtures including SYTO62 (Invitrogen, 1:500 dilution), anti-IgA (PE, eBioscience, clone:11-44-2, dilution 1:40) for IgA-seq samples, and for IgG-seq samples: anti- IgG1 PE (Beckton-Dickinson, #550083, 1:300) anti-IgG2ab PE (Beckton-Dickinson, #340269, 1:20), anti-IgG3 PE (Santa Cruz Biotech, #sc-3767, 1:1500). All stains were performed for 15 minutes at 4C. Stains were simultaneously performed using isotype control antibodies conjugated to PE, which yielded minimal to no staining for all experiments. Samples were subsequently washed and subjected to magnetic column-based fractionation as described above for human IgG- and IgA-seq. Bacterial DNA extraction and 16 rRNA amplification, pooling, and sequencing, were performed also as described above for human IgG- and IgA-seq.

16S rRNA sequence data were processed as described above for human IgA- and IgG-seq using Dada2 and rarefying sequences at 21,000 reads per sample. IgA and IgG scores were calculated as the log ratio of counts of each taxon in the Ig-enriched microbial fraction over counts in the Ig-depleted microbial fraction. Scores were compared across groups using non-parametric Mann-Whitney U tests.

### Design and construction of the human surrogate fecal community (SFC)

Stools from 40 healthy human donors were obtained (Lee BioSolutions, Maryland Heights, MO, USA) and subjected to DNA extraction and V4 16S rRNA amplification, purification, and pooling as described for paired IgA- and IgG-seq. Pooled libraries were sequenced on the NextSeq instrument (Illumina) using the custom R2 primer: “A+GTCAGTCAGCC+GGACTACHVGGGTWT+CTAAT” with locked nucleic acid base modifications denoted by “+”. PhiX was loaded at 40% concentration. ASV tables were generated using Dada2, and samples were rarefied to 140,000 sequences per sample. To ensure capture of IgG titers against the greatest number of human gut microbiota members, we sought to identify the combination of samples that would yield the most diverse final community as quantified by Shannon diversity. *In silico* modeling was performed by randomly combining 5, 10, 15, 20, 25, 30, 35, and all 40 samples 500,000 times each. Samples were concurrently assigned random proportions of the total *in silico-*mixed community during each permutation. The combination of 15 subjects yielding the maximum Shannon diversity was then chosen. Samples were resuspended in PBS with 1% bovine serum albumin (%w/v), mixed via gentle blending using a 21-ounce food processor, filtered through sterile 40 micron filters, OD-normalized, and added at the relative proportions dictated by the *in silico* modeling. The resultant SFC was centrifuged at 8,000 rcf for 3.5 minutes and resuspend in 15% glycerol in phosphate buffered saline at a concentration and volume to provide sufficient SFC for 96 wells per aliquot.

### IgG-seq using the SFC

An aliquot of the SFC was thawed, centrifuged, and washed with PBS with 1% bovine serum albumin (%w/v), a buffer solution used for all subsequent stain and wash steps. Human study participants for SFC-IgG-seq (IBD and healthy subjects, referring to data shown in Figures 3-5) provided informed consent and plasma was collected as part of a study approved by the Cedars-Sinai Medical Center Institutional Review Board. SFC aliquots were incubated in duplicate wells of 100 ul human subject plasma diluted to 1:300 for 30 minutes at 4C. An aliquot (1/100^th^ volume) was taken for flow cytometry-based assessment of IgG binding. Samples were then washed via centrifugation at 6,000 rcf for 4 minutes and resuspended in 20 ul of microbeads bound to IgG. Microbeads were made by conjugating reduced recombinant Protein G with N-terminal cysteine (Novus Biologicals) to amine functionalized magnetic beads (Alpha Biobeads) using LC-SMCC (succinimidyl 4-(N-maleimidomethyl)cyclohexane-1-carboxy-(6-amidocaproate)), (ThermoFisher) using the Alpha Biobeads manufacturer protocol. Plasma-treated SFC aliquots resuspended with protein G-microbeads were then shaken at 400 rpm for 20 minutes at room temperature. Samples were then placed on a 96-well plate magnet (Permagen, #MSP750) and allowed to bind for 2 minutes at room temperature. 30 ul of unbound suspension (IgG-unbound, ‘negative fraction’) were removed from these plates and a fixed quantity (5*10^6^ colony forming units) of *Staphylococcus hominis*, a human skin bacterium that was not present in the SFC, was added to each well to facilitate absolute quantification of taxa in the negative fraction. The suspension was frozen for downstream 16S rRNA amplification and sequencing on the NextSeq instrument (Illumina) as described above. A 1ul aliquot was taken from these negative fractions for flow cytometry-based quality check, in which cells were stained with anti-IgG PE (Miltenyi #130-093-193, 1:50 dilution) and SYTO-62 (Invitrogen, 1:500 dilution) for 20 minutes at 4C, along with the pre-enrichment and post-serum incubation aliquots taken at the start of the procedure. Samples were washed, fixed in 2% PFA for 15 minutes at 4C, and were acquired on a Fortessa cytometer (Beckton-Dickinson) with side-scatter thresholds set to 200. All QC assays for samples presented herein showed efficient depletion of IgG-bound bacteria in negative fractions with a range of 0.4-1.8% SFC bacteria remaining bound by IgG as compared to 4-29% bound in the pre-enrichment aliquots.

### SFC-IgG-seq analysis

Sequences were processed using Dada2(*64*) as above. Samples were rarefied at 400,000 reads per sample. All taxon abundances for each sample were divided by abundance of the *Staphylococcus hominis* control bacterium added for absolute quantification purposes, after which ASV abundances were given a pseudocount of 1 and log_10_-transformed. IgG scores were defined as log ratios of taxon abundance in the pre-enrichment SFC over their abundance in the negative fraction. Thus, ASV abundances in quadruplicate wells of the pre-enrichment SFC were averaged and ASV abundances in each sample were subtracted from the averaged quadruplicate pre-enrichment SFC ASV abundances. All resulting IgG scores were averaged across sample duplicates.

Bacterial taxa were determined to be bona fide translocators if they were cultured from the mesenteric adipose tissue of more than four human subjects in our cohort, and if an ASV with 100% identity at the 16S V4 rRNA region to the same bacterium by BLAST was detected in their gut mucosal microbiome as obtained and sequenced by Ha et al.(*34*) and previously deposited in BioProject: PRJNA659515. These 16S rRNA amplicon sequence data were processed via dada2 and each subject’s mucosal samples were aggregated by summation for the purpose of detecting presence of absence of translocator bacteria at the gut mucosa. Subjects with fewer than 19,000 total reads were excluded from analysis. Corresponding ASVs in the SFC were identified as those having >99% identity to the translocator bacterial species, and IgG scores for those taxa were compared between healthy healthy control subjects (who did not undergo mesenteric adipose tissue sampling and culture), IBD translocation-positive subjects (those for whom that bacterium was cultured from the mesenteric adipose tissue), and IBD translocation-negative subjects (who underwent mesenteric adipose tissue sampling but that bacterium was not cultured from their mesenteric samples). Linear regression was performed using the aforementioned categories as ordinal variables in the presented order, considering the mean row z-scores of IgG scores (centered and scaled IgG-scores per taxon across all subjects) for translocator taxa within each grouping, including all taxa as presented in Figure 3C. For individual translocator taxon IgG score comparisons, linear regression was performed using the same subject groupings as ordinal variables as above, and with row z-scores of IgG scores from each individual human subject, with resulting data presented in Figure 3D.

Comparisons of IgG scores between IBD subject groups (e.g. healthy, CD, and UC) were performed using non-parametric statistical tests in the base ‘stats’ library in R. For two-group comparisons, Mann-Whitney U tests were used (‘wilcox.test’), and for three-group comparisons, the Kruskal-Wallis test was used (‘kruskal.test’). For such analyses, ASVs that had non-zero count values in greater than 70% of samples were examined. Data were visualized using the package ‘superheat’ in R.

For visualization and comparison of total anti-microbiota IgG repertoires, an n-by-n matrix of pairwise comparisons was constructed using Pearson correlation coefficients comparing all IgG scores per sample to those of every other sample individually. Significance of group clustering was assessed using the resulting pairwise Pearson correlation matrices and the PERMANOVA technique (‘adonis’ function from package ‘vegan’) in R. Data were visualized using principal coordinates analyses also performed in R (‘pco’ from package ‘labdsv’).

### Bacterial activity and growth rate assessments in external validation datasets

Raw shotgun metagenomics reads from the Mayo Clinic study (*50*) were obtained from SRA (BioProject accession: PRJNA487636) and processed using MetaPhlAn2(*65*) to generate taxonomic relative abundance tables and GRiD(*48*) for quantification of peak-to-trough (PTR) ratios. Reads from the MetaHIT study(*49*) were obtained from the ENA database (accession: PRJEB1220) and processed for PTR ratios using GRiD, and using curatedMetagenomicData(*66*) to derive taxonomic relative abundances. The iHMP study data were obtained from Supplemental Tables 15 and 18 of the original publication(*45*). The continuous variable of dysbiosis was defined in the iHMP study in detail in the original publication. Briefly, dysbiosis scores were calculated for each individual as Bray-Curtis similarities to all healthy subject gut microbiota profiles, with higher scores denoting a greater deviation in microbiota community composition of healthy healthy subjects. All taxonomic relative abundance and GRiD values were compared between Crohn’s disease subjects and healthy controls for the Mayo Clinic and MetaHIT studies using non-parametric Mann-Whitney U tests.

To minimize the effects of microbiota-associated confounding variables(*1*), CD and healthy sample populations were matched for body mass index, age, and sex where data were available. For the MetaHIT study by Nielsen et al., the ‘healthy’ subject group was chosen as this group was matched for age and sex to the CD group (age P=0.93 by two-sided T test, sex P=0.62 by chi-square test), while the ‘healthy relatives’ group was not matched for age (P= 3.9*10^-5^ by two-sided T test) to the CD group. This yielded a study population of 13 CD subjects and 10 healthy control subjects. For the Mayo clinic study by Muñiz Pedrogo et al., subjects with rheumatoid arthritis or arthropathy were excluded from both the CD group and healthy control groups. Inclusion criteria were imposed to ensure matching for age (>25 years old) and BMI (<40 and >20 BMI), which yielded a well-matched study population for age (P=0.54 by two-sided T test), sex (P=0.56 by chi-square test), and BMI (P=0.42 by two-sided T test) of N=63 controls and N=32 CD subjects.

To facilitate comparisons to our IBD IgG-seq and translocator bacteria dataset, analyses of relative abundances and bacterial growth rate were limited to taxa classified at the species level in the metagenomics datasets. Taxa were considered translocators if they were cultured from the mesenteric adipose tissue of >4 subjects in the study by Ha et al.(*34*). Taxa were considered as having significantly higher IgG scores in CD subjects if they passed the following criteria: 1) unadjusted P<0.05 in Mann-Whitney U tests comparing CD to healthy subjects. 2) unadjusted P<0.02 in Kruskal-Wallis tests comparing CD, UC, and healthy subjects as in the analyses of Figure 4. 3) Greater mean IgG score in CD compared to healthy subjects. 4) >99% v4 16S rRNA sequence identity to its nearest species classification by BLAST.

## Acknowledgments

We wish to thank Apollo Stacey (NIH/NIAID), Naisarg Modi (NIH/NIAID), Seong-Ji Han (NIH/NIAID), Shurjo Sen (NIH/NCI), and Soo Ching Lee (NIH/NIAID) for assistance and helpful discussion. We also thank Les Laboratoires Servier (https://smart.servier.com/). We thank Dr. Michael Grigg (NIAID/NIH) for sharing an RFP-encoding *T. gondii* strain. We wish to thank the CEPHIA consortium and its funding sources, including the Bill and Melinda Gates Foundation (OPP1017716, OPP1062806 and OPP1115799), the National Institutes of Health (P01 AI071713, R01 HD074511, P30 AI027763, R24 AI067039, U01 AI043638, P01 AI074621 and R24 AI106039); the HIV Prevention Trials Network (HPTN) sponsored by the NIAID, National Institutes of Child Health and Human Development (NICH/HD), National Institute on Drug Abuse, National Institute of Mental Health, and Office of AIDS Research of the NIH, DHHS (UM1 AI068613 and R01 AI095068); the California HIV-1 Research Program (RN07-SD-702); Brazilian Program for STD and AIDS, Ministry of Health (914/BRA/3014-UNESCO); and the São Paulo City Health Department (2004–0.168.922– 7). The content of this publication does not necessarily reflect the views or policies of DHHS, nor does the mention of trade names, commercial products, or organizations imply endorsement by the U.S. Government.

## Author contributions

Conceptualization: IVC, YB

Methodology: IVC, HW, CHWH, JMB

Formal analysis: IVC

Investigation: IVC, SM, CHWH, HW, LH

Visualization: IVC

Funding acquisition: YB, SD

Project administration: YB

Writing – original draft: IVC

Writing – review & editing: YB, IVC

Final approval of manuscript: All authors

## Supplemental Figures

**Figure S1:**
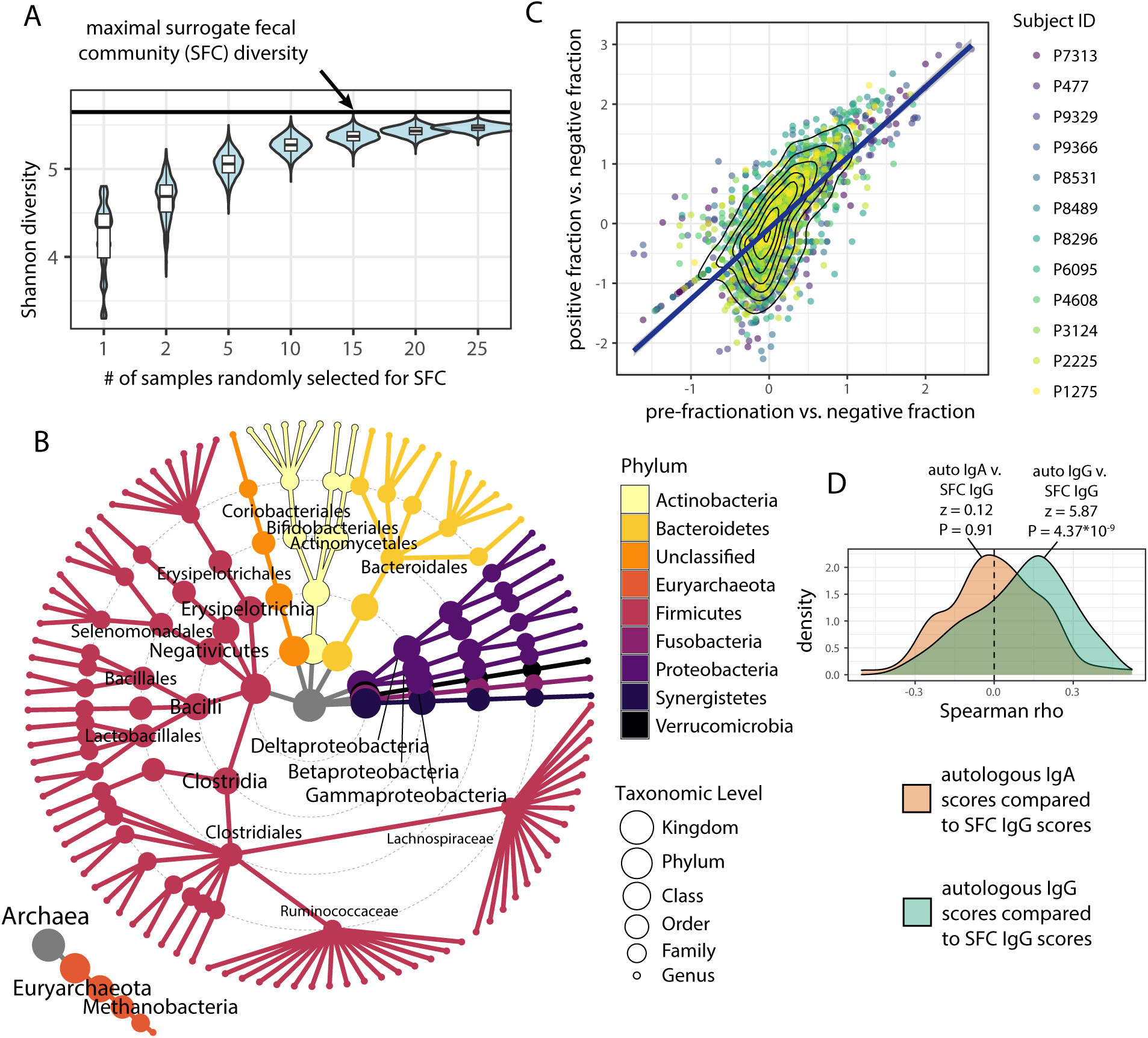
Design and validation experiments pertaining to the surrogate fecal community (SFC). A) As described in Methods, forty healthy human fecal donor samples were profiled via 16S rRNA sequencing. Random permutations of combinations of human subject fecal communities were assessed for overall diversity as calculated by the Shannon diversity index, in combinations of 1, 2, 5, 10, 15, 20, and 25 subjects at a time. A plateau for maximal SFC diversity (shown as solid black line) was observed beginning at combinations of 15 subjects, and the maximum diversity combination of 15 subjects was chosen for physical construction of the SFC. B) Taxonomic distribution of amplicon sequence variants robustly detected in SFC, with phyla denoted by color. Taxa were considered present if their relative abundance was >0.001% of total sequences and were detected in >70% of samples. C) Validation of usage of the pre-enrichment fraction in place of the IgG-bound (“positive”) fraction for calculating IgG scores. Data generated as part of Figure 1 in the main text were utilized to test whether high-throughput profiling via the SFC could be streamlined via comparison of IgG-unbound fractions to the pre-enrichment SFC instead of to IgG-bound fractions as was performed in data for Figure 1. A strong concordance was seen between the two methods (Spearman correlation P<10^-16^). D) The subject samples profiled for IgA and IgG scores using autologous stool and plasma were subjected to SFC-IgG-seq using the SFC in place of autologous stool for comparison of the two methods. IgG scores from the SFC-IgG-seq method had correlations with IgA scores from the autologous method that were consistent with randomness. IgG scores from the two methods compared to each other had non-random concordance (two-sided Z-test P=4.37*10^-9^).

**Figure S2:**
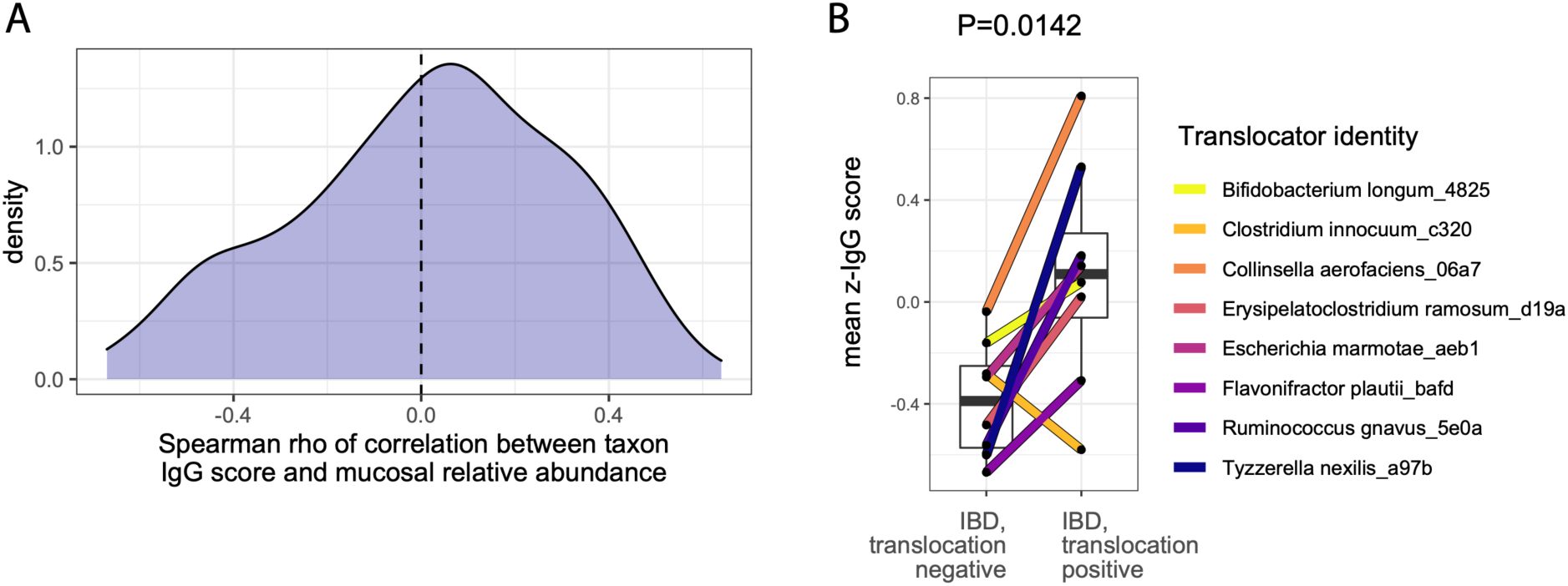
Comparison of IgG scores to mucosal relative abundances in IBD subjects. Gut mucosal tissue samples taken at time of surgical resection were subjected to 16S rRNA profiling as described in Ha et al. (*2*) and data were obtained from PRJNA659515. A) Spearman correlation tests comparing mucosal relative abundance and IgG score for each ASV were performed. Distribution of resulting rho correlation coefficients is shown, and is consistent with a random distribution (Z-test P=0.478). B) A similar analysis was performed as is shown in Fig. 3C, limited to IBD subjects (no healthy controls) and only subjects for whom the translocator had detectable abundance in their mucosal 16S profiles. Bacteria exhibited heightened IgG scores in subjects with confirmed translocation as compared to those without translocation (P=0.0142, paired two-sided Student’s T-test).

**Figure S3:**
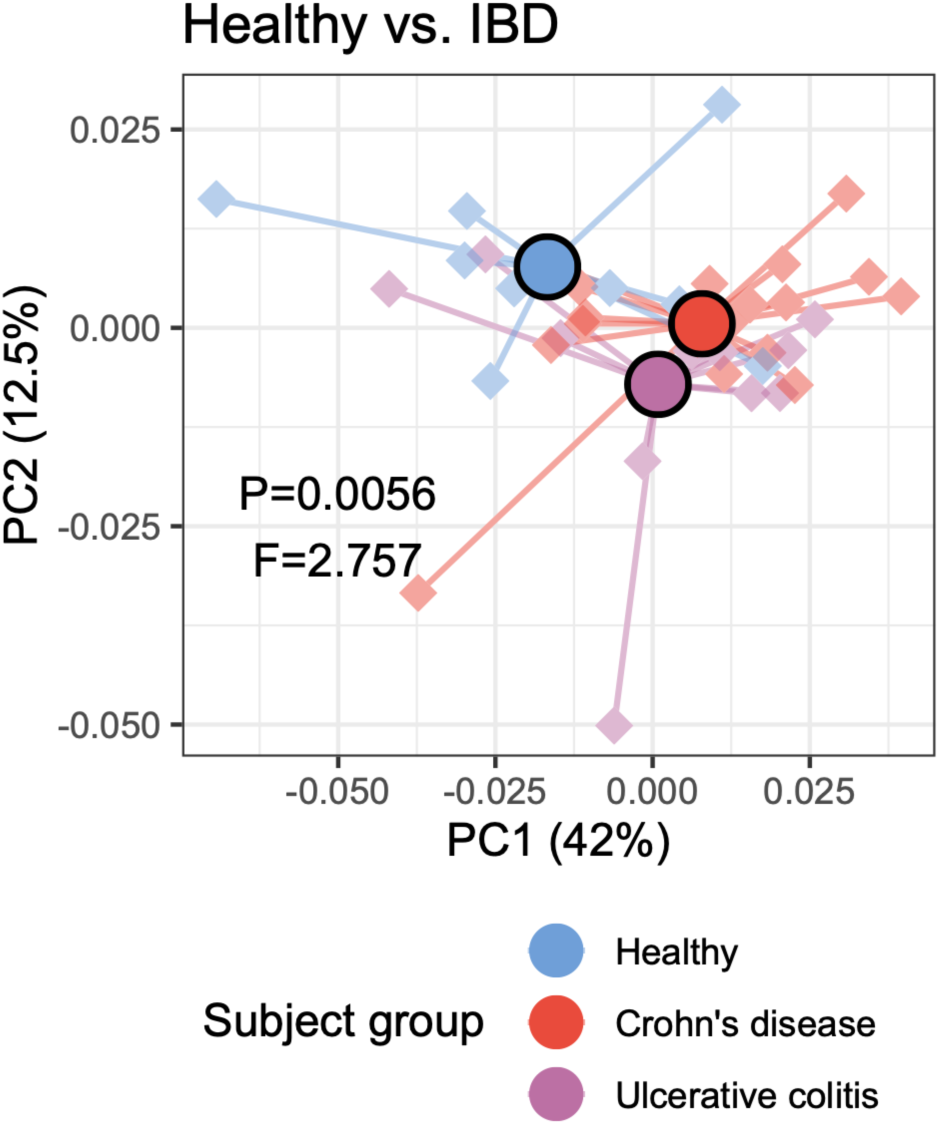
Principal coordinates analyses using pairwise Pearson correlation coefficient matrices encompassing IgG scores for all taxa, comparing both IBD subgroups together to healthy healthy subjects. Significance of clustering was tested by PERMANOVA (P=0.0056).

